# A Novel Type 2 Diabetes Locus in sub-Saharan Africans, *ZRANB3*, is Implicated in Beta Cell Proliferation

**DOI:** 10.1101/646513

**Authors:** Adebowale A. Adeyemo, Guanjie Chen, Ayo P. Doumatey, Timothy L. Hostelley, Carmen C. Leitch, Jie Zhou, Amy R. Bentley, Daniel Shriner, Olufemi Fasanmade, Godfrey Okafor, Benjamin Eghan, Kofi Agyenim-Boateng, Settara Chandrasekharappa, Jokotade Adeleye, William Balogun, Samuel Owusu, Albert Amoah, Joseph Acheampong, Thomas Johnson, Johnnie Oli, Clement Adebamowo, South Africa Zulu Type 2 Diabetes Case-Control Study, Francis Collins, Georgia Dunston, Norann A. Zaghloul, Charles N. Rotimi

## Abstract

Genome analysis of diverse human populations has contributed to the identification of novel genomic loci for diseases of major clinical and public health impact. Here, we report the largest genome-wide analysis of type 2 diabetes (T2D) in sub-Saharan Africans, an understudied ancestral group. We analyzed ~18 million autosomal SNPs in 5,231 individuals from Nigeria, Ghana and Kenya. *TCF7L2* rs7903156 was the most significant locus (p=7.288 × 10^−13^). We identified a novel genome wide significant locus: *ZRANB3* (Zinc Finger RANBP2-Type Containing 3, lead SNP chr2:136064024, T allele frequency=0.034, p=2.831×10^−9^). Knockdown of the zebrafish ortholog resulted in reduction in pancreatic beta cell number in the developing organism, suggesting a potential mechanism for its effect on glucose hemostasis. We also showed transferability in our study of 32 established T2D loci. Our findings provide evidence of a novel candidate T2D locus and advance understanding of the genetics of T2D in non-European ancestry populations.

---

The genetic architecture of type 2 diabetes (T2D, MIM:125853) in Africa remains largely understudied. While the reduced linkage disequilibrium (LD) characteristic of African populations was used to refine and fine map the original *TCF7L2* genetic association^1,2^, genome-wide and/or high throughput studies of the genetics of T2D in Africa remain limited to a genome-wide linkage analysis^3^ and a large-scale replication study^4^, both from the Africa America Diabetes Mellitus (AADM) Study. African American populations, on the other hand, have been studied more comprehensively including several genome-wide association studies (GWAS) and meta-analysis of GWAS^5^. However, African American populations should not be used as proxies for populations in Africa because of differences in genetic (African Americans have ~ 20% European admixture) as well as non-genetic risk factors (including lifestyle and behavioral factors). Therefore, despite the advances over the last decade in our understanding of the role of genetic variants influencing T2D risk and the identification of the role of the genes in pathophysiology, data from Africa remains scarce.

In the present study, we conducted a GWAS of T2D in Africa using data from over 5,000 Africans enrolled from Nigeria, Ghana and Kenya as part of the Africa America Diabetes Mellitus (AADM) study^3,6^ and extend the transferability of previously reported T2D loci in Africa. We identified a novel genome-wide significant locus for T2D – the *Zinc Finger RANBP2-Type Containing 3 (ZRANB3)* gene. Functional studies of the *ZRANB3* ortholog in zebrafish showed that knockdown of the gene leads to reduction in beta cell number in the developing embryo. Our findings represent an advance in our knowledge of the genetics of T2D in sub-Saharan Africa.

## Results

### Characteristics of discovery sample

The characteristics of the 5231 AADM study participants (2,342 T2D cases and 2,889 controls) are shown in Table 1. T2D cases were older than controls (mean age 55 years versus 46 years). Mean BMI was similar between cases and controls. However, cases had a significantly bigger waist circumference than controls (mean 93.7 cm versus 88.5 cm). Fasting glucose values indicate that most of the T2D cases were uncontrolled at the time of enrolment with a median fasting glucose of 153 mg/dl (8.5 mmol/L) and more than three-quarters having fasting glucose values greater than 109 mg/dl (6.1 mmol/L) at the time of enrollment into the study.

**Table 1:**
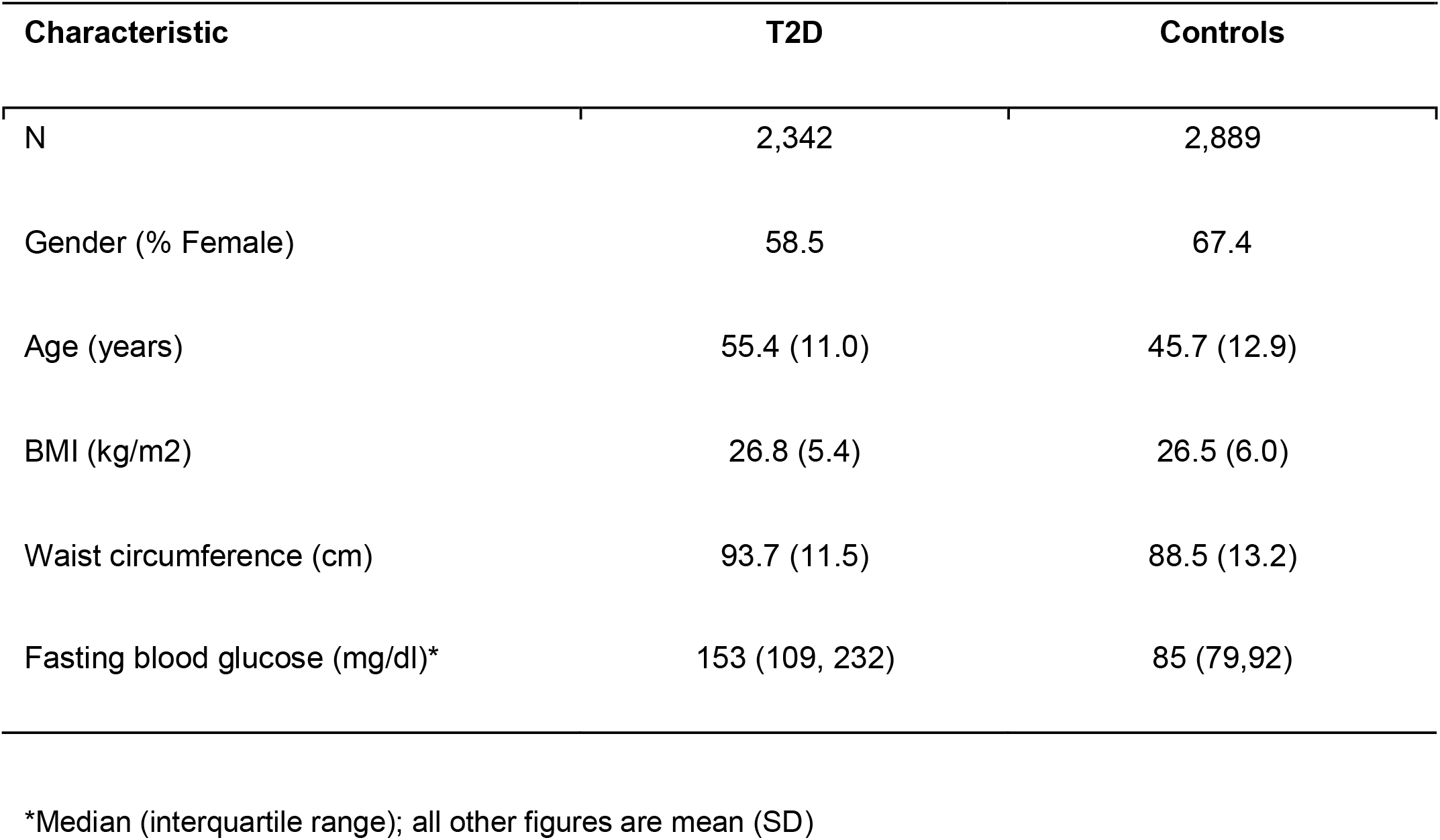
Characteristics of discovery study sample in the AADM Study

### Discovery genetic association analysis

The distribution of association statistics for the genome-wide association analysis is shown in the Manhattan plot (Figure 1). There was minimal inflation of the association statistics (λ = 1.013, Supplementary Figure S2). Three genome-wide significant loci were identified (Table 2): *TCF7L2* (lead SNP rs7903146, T allele frequency=0.331, p=7.288 × 10^−13^), *HMGA2* (lead marker rs138066904, deletion frequency=0.096, p=2.516 × 10^−9^) and *ZRANB3* - Zinc Finger RANBP2-Type Containing 3 (lead SNP chr2:136064024, T allele frequency=0.034, p=2.831×10^−9^). *TCF7L2* is an established T2D risk locus and the lead SNP of *TCF7L2* (rs7903146) in the present study is the same lead SNP reported in most GWAS of T2D to date (Figure 2). *HMGA2* is also a known T2D locus in both Europeans and African Americans. However, the genome-wide significant *HMGA2* variant in the present study is a deletion (CCTAG/C), not a SNP like other *HMGA2* markers that have previously been found to be genome-wide significant for T2D in Europeans (leading SNP rs2258238, 68.5 kb away from the deletion) and in African Americans (leading SNP rs343092, 38.6 kb away). The LD between the deletion and these other SNPs is low (r^2^ 0.052 and 0.003, respectively) in this study of sub-Saharan Africans.

**Figure 1:**
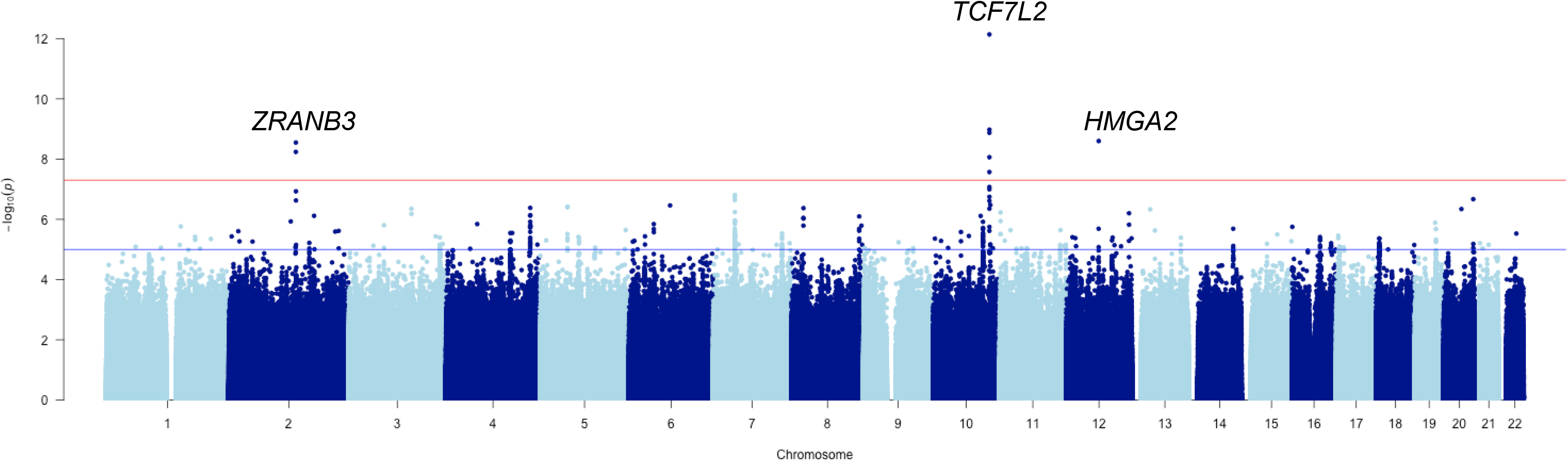
Manhattan plot of Discovery GWAS: the AADM Study

**Figure 2:**
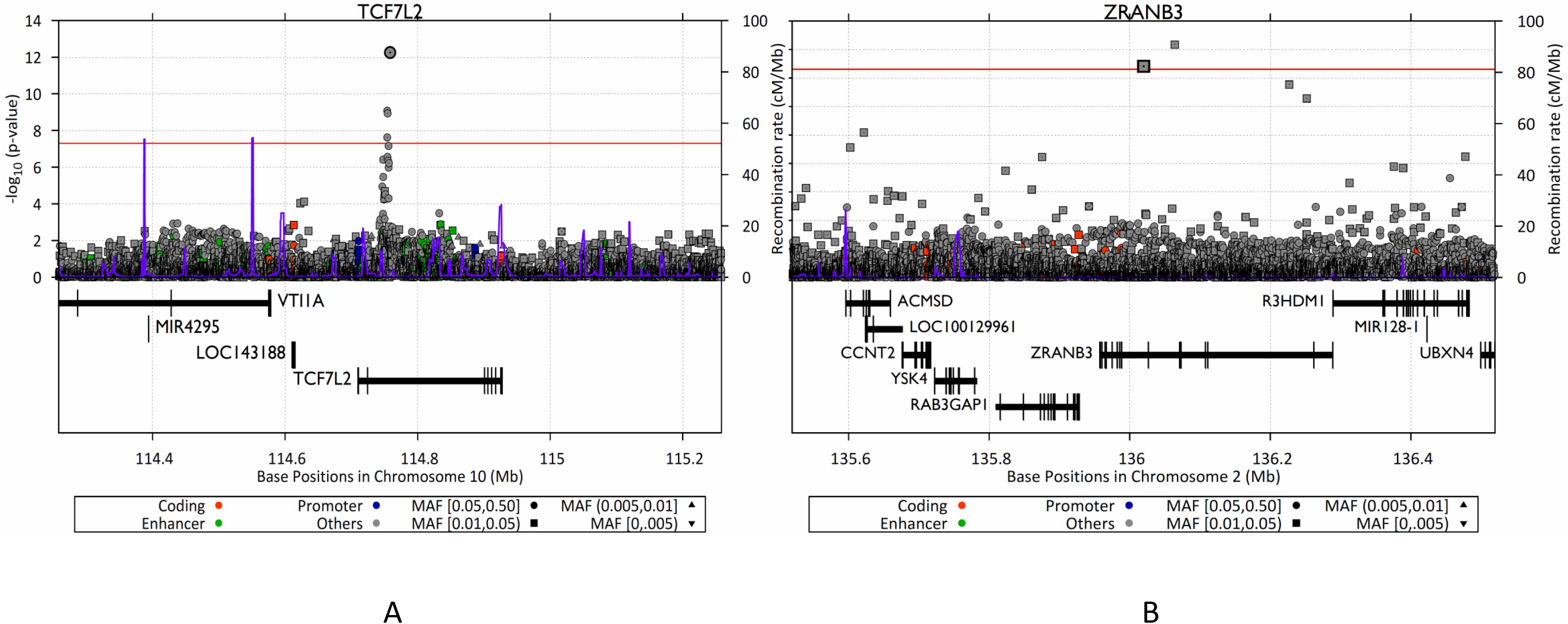
Regional association plots for *TCF7L2* and *ZRANB3* in the AADM GWAS for T2D

**Table 2:**
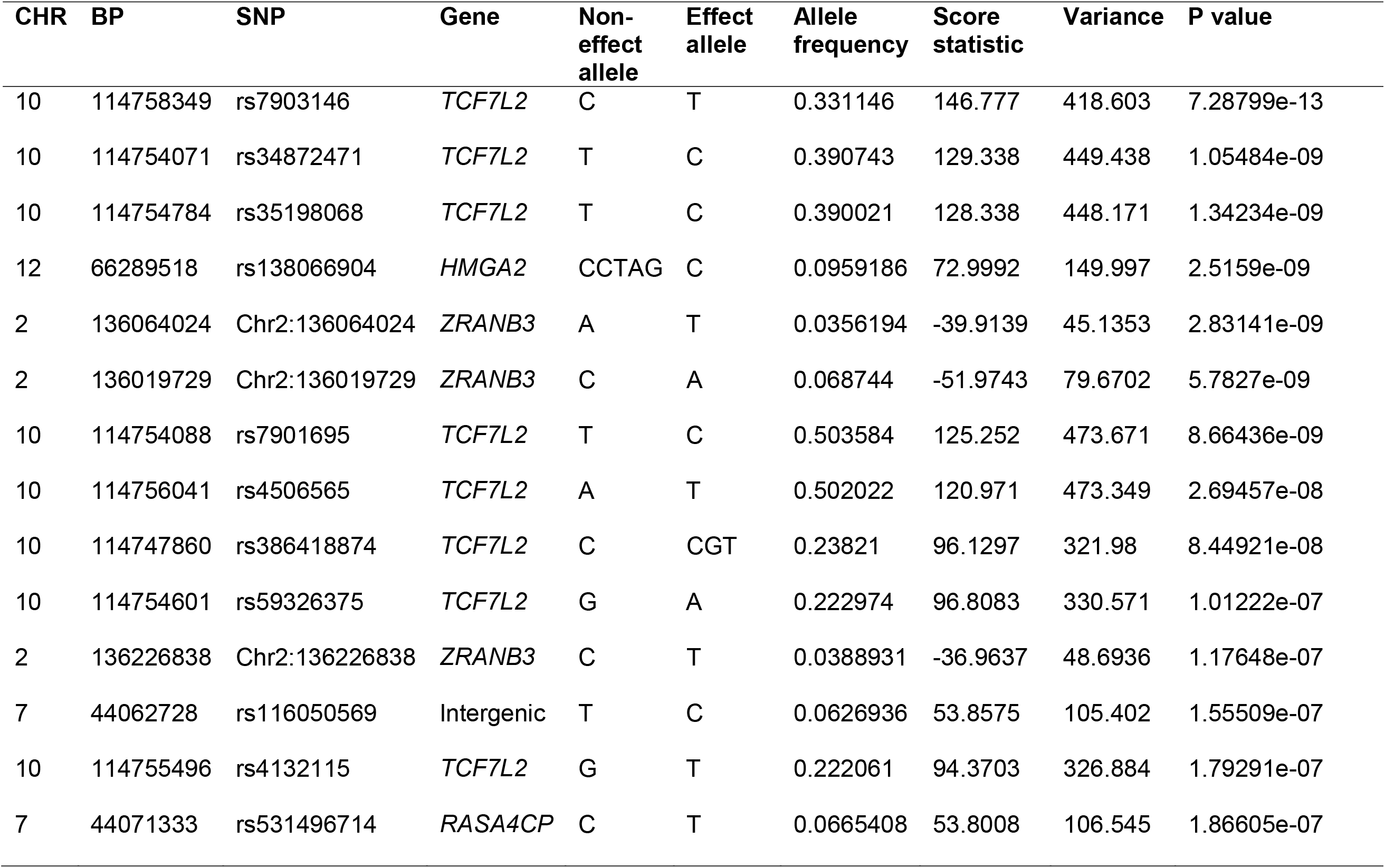
SNPs most significantly associated with T2D in 5,231 sub-Saharan Africans (the AADM Study)

### Replication and Annotation of ZRANB3

*ZRANB3* is a novel candidate locus for T2D as it has not been previously reported in relation to T2D. The leading SNPs have a frequency of 3.6% and 6.9%, respectively (Table 3). Of the top four markers in *ZRANB3*, three appear to be African-specific as they are not present in other populations as evaluated through the 1000 Genomes and gNOMAD databases. One marker (ZRANB3 2:136019729, rs755264566) is present in the gNOMAD database with an overall frequency of 5.5%. For evaluation of replication of ZRANB3 in another African ancestry population, we examined these variants in South African Zulu T2D cases and controls from the Durban Diabetes Case-Control Study (DCC) and the Durban Diabetes Study (DDS) - Table 3. The leading SNPs in *ZRANB3* in AADM each showed consistency of direction of effect in the Zulu GWAS, despite the latter study showing lower effect allele frequencies (chr2:136064024: T allele frequency 0.9% Zulu versus 3.4% AADM; chr2:136019729 A allele frequency 2.6% Zulu versus 6.9% in AADM). The combined p-values for the two leading SNPs across the discovery and replication samples remained genome-wide significant (Table 3).

**Table 3:** Association tests for the *ZRANB3* T2D locus in AADM discovery (n=5,231) and Zulu replication (n=2,578) studies

| Marker | EA | NEA | AADM |  |  |  | Zulu |  |  |  | Meta-analysis |  |  |
| --- | --- | --- | --- | --- | --- | --- | --- | --- | --- | --- | --- | --- | --- |
|  |  |  | EAF | BETA | SE | P value | EAF | BETA | SE | P value | Direction of effect | Z | P value |
| 2:136064024 | T | A | 0.036 | -0.8843 | 0.1488 | 2.83E-09 | 0.009 | -0.1565 | 0.1325 | 2.38E-01 | -- | -5.541 | 3.015E-08 |
| 2:136019729 | A | C | 0.069 | -0.6523 | 0.1120 | 5.78E-09 | 0.026 | -0.1022 | 0.05787 | 7.74E-02 | -- | -5.781 | 7.446E-09 |
| 2:136226838 | T | C | 0.039 | -0.7591 | 0.1433 | 1.18E-07 | 0.014 | -0.03315 | 0.08541 | 6.97E-01 | -- | -4.559 | 5.147E-06 |
| 2:136251877 | A | C | 0.076 | -0.6063 | 0.1173 | 2.36E-07 | 0.047 | -0.01784 | 0.03502 | 6.10E-01 | -- | -4.523 | 6.090E-06 |

*ZRANB3* is a protein-coding gene with nucleic acid binding and endonuclease activity. The *ZRANB3* transcript is the target of nonsense mediated decay (NMD) and is expressed in tissues relevant to T2D including adipose tissue, skeletal muscle, pancreas and liver (Supplementary Figure S3). We identified haplotype blocks around the two genome-wide significant *ZRANB3* SNPs and identified 35 and 43 target genes, respectively, from the significant t*rans*-eQTL-gene associations in each haplotype block using the Framingham Heart Study (FHS) eQTL database (Table 4). We also identified five common target genes for cis-eQTLs in addition to *ZRANB3*. Overlaying these associations with known T2D loci from the GWAS Catalog highlighted two known-T2D genes associated with *cis*-eQTLs (*MCM6, DARS*) and four with *trans*-eQTLs (*DGKB, GTF3AP5-AGMO, IL23R/IL12RB2, SLC44A4*). It is noteworthy that there is a ClinVar record of a duplication in *ZRANB3* associated with gestational diabetes (ClinVar Accession SCV000191187.1: see **Web Resources**). Also, data from the Rat Genome Database shows that the syntenic region contains QTLs for insulin levels (insulin level QTL 44) and glucose level (glucose level QTLs 66 and 67) in the rat. At the present time (October 2017), there is no *ZRANB3* variant in the NHGRI-EBI GWAS Catalog or in OMIM

**Table 4:** eQTL* annotation of ZRANB3 genome wide significant SNPs in the AADM Study

| Index SNP | Haplotype block** |  |  | Flanking SNPs |  |  |  |
| --- | --- | --- | --- | --- | --- | --- | --- |
|  | *hg19*<br>coordinates<br>(size) | Number of<br>target genes<br>for which<br>eQTLs<br>Is in *cis/trans* | eQTL for T2D-related genes<br>in GWAS<br>Catalog | SNP ID | Is *cis*-eQTL<br>for *ZRANB3* | Is *trans*-<br>eQTL<br>(genes) | eQTL for<br>T2D-related<br>genes in<br>GWAS<br>Catalog |
| 2:136064024 | 2:136046052-136064758<br>(18.7 kb) | 5/43 | *MCM6*<br>*DARS*<br>*IL23R/IL12RB2*<br>*SLC44A4*<br>*ZFAT* | rs57510176 | Yes | Yes<br>*ATXN10* | - |
|  |  |  |  | rs72982351 | Yes | No | - |
| 2:136019729 | 2:136016079-136032905<br>(16.8 kb) | 5/35 | *MCM6*<br>*DARS*<br>*DGKB*<br>*GTF3AP5-*<br>*AGMO*<br>*IL23R/IL12RB2*<br>*SLC44A4* | rs141414859 | Yes | Yes<br>*DGKB*,<br>*CGREF1* | *DGKB* |
|  |  |  |  | rs112334908 | Yes | No |  |

\*eQTL data from the Framingham Heart Study (FHS) eQTL database<sup>11</sup> accessed through the NCBI Molecular QTL Browser
\*\*Haplotype block around index SNP defined using 1000 Genomes Phase 3 AFR data

### Replication of known T2D loci

We investigated the transferability of previously reported T2D SNPs in the AADM sample. Of the 130 SNPs, 108 were present in our dataset. Sixteen SNPs (or 15%) showed exact replication, i.e. consistent direction of effect for the alleles and p < 0.05 – Table 5. Two other SNPs [rs2258238 (*HMGA2*) and rs12595616 (*PRC1*)] showed a p < 0.05 with inconsistent direction of effect. Sixteen other loci showed local replication, including *KCNJ11, HHEX/IDE, THADA, MC4R* and *ATP8B2* (Supplementary Table S1). Of the two loci first reported in an African ancestry GWAS meta-analysis for T2D (the MEDIA Consortium), we found exact replication for *INS-IGF2* rs3842770 (p=1.867 × 10^−3^, Table 4) but no evidence for replication for *HLA-B* rs2244020 (rs74995800, p=0.798).

**Table 5:** Exact replication of established T2D loci in 5231 Sub-Saharan Africans

| SNP | CHR | BP | EA | NEA | EAF | BETA | SE | P Value | Locus |
| --- | --- | --- | --- | --- | --- | --- | --- | --- | --- |
| rs7903146 | 10 | 114758349 | T | C | 0.331 | 0.350635 | 0.0488764 | 7.29E-13 | *TCF7L2* |
| rs10998572 | 10 | 70859204 | A | C | 0.055 | -0.244687 | 0.104606 | 1.933E-02 | *VPS26A* |
| rs231360 | 11 | 2692249 | T | C | 0.620 | 0.145475 | 0.047877 | 2.378E-03 | *KCNQ1* |
| rs441613 | 11 | 2910191 | C | T | 0.605 | 0.115247 | 0.0471471 | 1.451E-02 | *KCNQ1* |
| rs2258238 | 12 | 66221060 | T | A | 0.367 | -0.100573 | 0.048316 | 3.738E-02 | *HMGA2* |
| rs12595616 | 15 | 91563513 | C | T | 0.852 | -0.227137 | 0.0656811 | 5.438E-04 | *PRC1* |
| rs1558902 | 16 | 53803574 | A | T | 0.050 | 0.244757 | 0.104983 | 1.973E-02 | *FTO* |
| rs2925979 | 16 | 81534790 | C | T | 0.676 | -0.106593 | 0.0489676 | 2.950E-02 | *CMIP* |
| rs7224685 | 17 | 4014384 | T | G | 0.435 | 0.147026 | 0.0460284 | 1.402E-03 | *ZZEF1* |
| rs13342692 | 17 | 6946287 | C | T | 0.393 | 0.145304 | 0.0470417 | 2.009E-03 | *SLC16A11/A13* |
| rs7234111 | 18 | 7067652 | C | T | 0.584 | 0.0947849 | 0.0471346 | 4.433E-02 | *LAMA1* |
| rs4402960 | 3 | 185511687 | T | G | 0.570 | 0.115527 | 0.0468621 | 1.369E-02 | *IGF2BP2* |
| rs1182436 | 7 | 157027753 | C | T | 0.326 | 0.10368 | 0.0499157 | 3.779E-02 | *MNX1* |
| rs3802177 | 8 | 118185025 | A | G | 0.038 | -0.310673 | 0.118332 | 8.654E-03 | *SLC30A8* |
| rs10757282 | 9 | 22133984 | C | T | 0.207 | 0.149237 | 0.0556943 | 7.372E-03 | *CDKN2A/B* |
| rs1575972 | 9 | 22301092 | A | T | 0.320 | -0.196379 | 0.0491555 | 6.47E-05 | *DMRTA1* |
| rs10758593 | 9 | 4292083 | A | G | 0.512 | 0.117341 | 0.0462949 | 1.126E-02 | *GLIS3* |
| rs3842770 | 11 | 2178670 | A | G | 0.291 | 0.124185 | 0.0527965 | 1.867E-02 | *INS-IGF2* |

### Meta-analysis with African ancestry populations and trans-ethnic meta-analysis

We conducted an African ancestry T2D meta-analysis that included this GWAS and an African American meta GWAS^7^ consisting of five studies (n=8599; the Atherosclerosis Risk in Communities (ARIC), the Cleveland Family Study (CFS), the Howard University Family Study (HUFS), Jackson Heart Study (JHS) and Multi-Ethnic Study of Atherosclerosis (MESA). This African ancestry meta-analysis revealed four genome-wide significant loci (Supplementary Figure S4, Supplementary Table S2), namely: *TCF7L2* (lead SNP rs386418874 p= 6.33 x 10^−11^), *TH-INS* (lead SNP rs4072825 p= 2.35X10^−8^), *KCNQ3* (lead SNP rs111248619 p= 2.71×10^−8^) and an intergenic locus (lead SNP rs4532315 p= 9.80×10^−9^).

Trans-ethnic meta-analysis of the above African ancestry studies with a large GWAS of European ancestry individuals (the DIAGRAM meta-analysis of type 2 diabetes (T2D) based on the GoT2D integrated haplotypes)^8^ revealed multiple genome-wide significant loci as expected. However, they are all known loci and none is a novel locus for T2D (data not shown).

### Suppression of zranb3 in zebrafish results in reduced β-cell mass

To examine a potential role for *zranb3* in T2D etiology *in vivo* we targeted the gene in transgenic zebrafish larvae in which β-cells could be visualized and quantified, Tg(*insa:mCherry*)^9^. Transgenic embryos were injected at the one-to two-cell stage with a morpholino (MO) designed to disrupt splicing of the endogenous *zranb3* transcript at exon 5. We first validated the efficacy of the MO to significantly suppress *zranb3* mRNA expression without inducing off-target toxicity by assessing transcript levels of endogenous *zranb3* and by examining the presence of a marker of MO-induced toxicity, the delta113 isoform of *p53*,^10^ respectively (Supplementary Figure S5). We cultured injected embryos to 5 days post fertilization (dpf) when the zebrafish principal islet is fully formed and functional and we could quantify β-cell number. Suppression of *zranb3* expression resulted in a significant reduction of β-cells produced (26.8 β-cells per larva compared with 33.3 in control larvae, p=0.0002). We confirmed that this effect was directly relevant to MO-induced knockdown as the impact on β-cell number increased with increased MO dose (Figure 3). To further validate the impact of disruption of *zranb3* on production of β-cells, we generated a model of genomic disruption of *zranb3* by CRISPR/Cas9-mediated targeting of either exon 5 or exon 7 of *zranb3*. Tg (*insa:mCherry*) embryos co-injected with either *zranb3*-targeted sgRNA and Cas9 mRNA were cultured until 5 dpf and β-cell numbers quantified. We observed a similar significant reduction of β-cell number with either sgRNA (exon 5: 28.5 compared with 33.5 in controls, p=0.0006; exon 7: 27.1 compared with 33.3 in controls, p=0.0001). The *zranb3* knockdown model exhibited an increase in ER stress as evidenced by qRT-PCR data showing increased expression for the ER stress marker genes BiP, CHOP and EDEM (Supplementary Figure S6).

**Figure 3:**
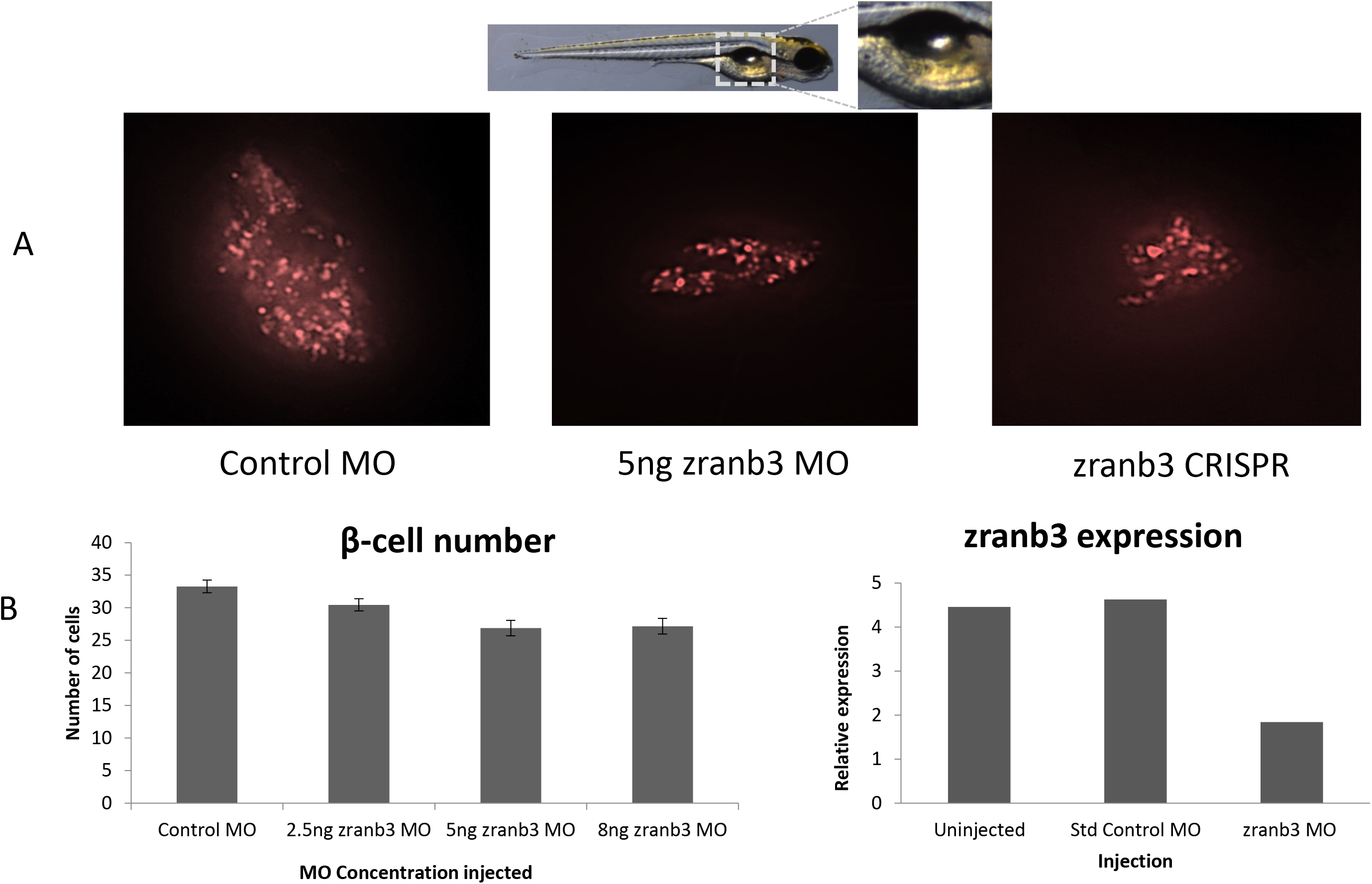
Effect of *ZRANB3* knockdown in zebrafish *Legend:* (A) mCherry expression in zebrafish β-cells at 5 days post fertilization (dpf) (B) Quantification of β-cell number in 5 dpf zebrafish (left panel) and Validation of zranb3 knockdown on zranb3 expression (right panel)

### Integrative analysis of AADM discovery GWAS with transcriptomic data

Integrative analysis that combines GWAS summary statistics with eQTL data was performed to identify potential new candidate genes. The most significant genes are shown in Figure 4. Nearly all of these genes are not known T2D risk loci with the exception of *MARCH1* which has been reported to be associated with T2D^11^. *LIPC* which is associated with closely related phenotypes, including the metabolic syndrome^12^ and circulating levels of total cholesterol, HDL-cholesterol and triglycerides in multiple studies.^13–19^

**Figure 4:**
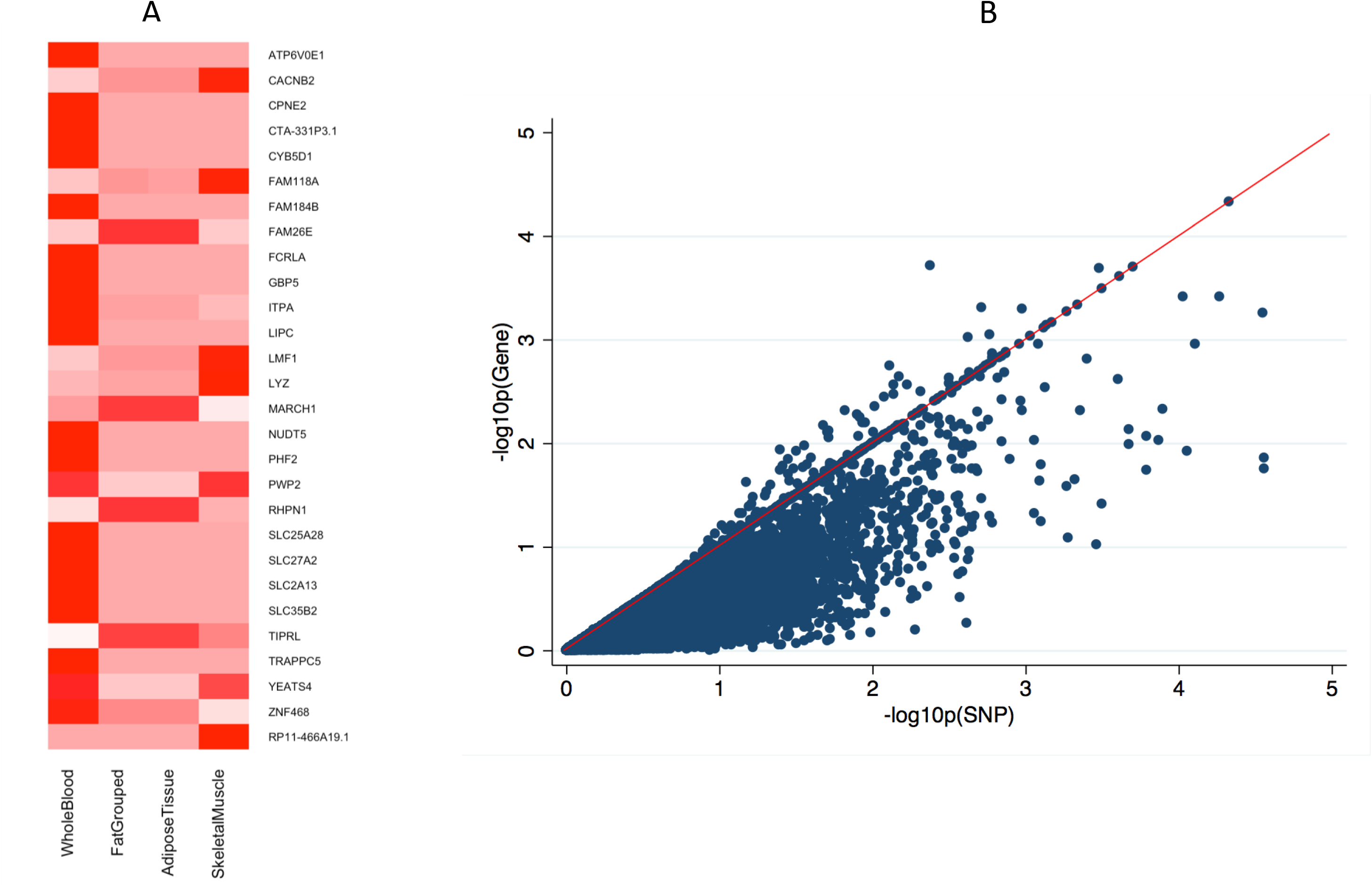
Significant genes on integrative GWAS and transcriptomic analysis. *Legend:* Panel A: Figure showing genes with gene-set p < 10^−3^ colored in deep red. Transcriptomic data from “Whole Blood” (11 studies), “Fat Grouped” (Grundberg et al, Nat Genet 2012 + GTeX adipose tissue), “Adipose Tissue” (GTeX adipose tissue only), “Skeletal Muscle” (GTeX skeletal muscle only). Panel B: Plot of gene-based versus best single variant association p-values for “Whole Blood”.

## Discussion

The vast diversity of genetic characteristics and environments across the world indicates that common complex disorders such as T2D need to be studied in diverse global populations. Nowhere is this truer than in sub-Saharan Africa, which is not only the cradle of humanity but is also home to a vast diversity of populations with widely divergent lifestyle, behavioral and environmental factors including long term exposure to pathogens that have shaped the genomic architecture of African peoples. In the present study, we report a genome-wide analysis of T2D in over 5,000 sub-Saharan Africans representing the largest sample size from a single diabetes association study from the continent to date. Reassuringly, *TCF7L2* rs7903146 was genome-wide significant as expected from previous T2D studies in Africa^4,20–25^ and consistent with the findings of most populations studied around the world. We also replicated several previously reported T2D loci. Using exact replication strategies (same SNP, consistent direction of effect, p value < 0.05), we demonstrated that 15% of the reported loci were significantly associated in this study of sub-Saharan Africans. This is consistent with our estimates of the power of our study to replicate known loci considering effect allele frequencies and reported effect sizes. Using local replication strategies, which help to identify significant loci which would otherwise have been missed because of allele frequency and/or LD differences between populations, we replicated additional sixteen loci (i.e. significant p value in a SNP in LD with the original reported SNP). These findings extend our previous studies of transferability of T2D loci in sub-Sharan Africa.^4^ Notably, the present study’s exact replication rate of 15% is marginally more than the 11% that we previously reported in a smaller sample from the same study. While this difference is not statistically significant (p=0.369, test of difference between proportions), it suggests that larger sample sizes may lead to increased numbers of replicated variants because of increased statistical power. It should be noted that transferability of reported T2D genome-wide significant variants between populations has always been demonstrable for a relatively small fraction of all such loci, especially with African ancestry populations.^26,27^. Perhaps this is not unexpected given that transferability is affected by several factors including sample sizes, effect allele frequencies, LD structure and genetic architecture of the trait. While acknowledging that increased sample size is just one of these variables, an appreciation of the final set of consistently replicable variants across populations will probably become possible when sample sizes (and the resultant statistical power) in non-European studies begin to approach those of European GWAS.

A notable finding in the present study is the identification of a promising novel locus (*ZRANB3*: Zinc Finger RANBP2-Type Containing 3) for T2D. Two intronic SNPs in the gene were genome-wide significant and the direction of effect was consistent for the top four SNPs in a South African Zulu sample with the meta-analysis p values of the two SNPs remaining genome-wide significant. The SNPs are African-specific and were discovered by sequencing of African genomes in the African Genome Resources Haplotype Reference Panel. Our functional annotation of the *ZRANB3* locus identified several *cis*- and *trans*-eQTLs, indicating that the locus contains multiple functional variants. Of relevance to T2D is our finding two known-T2D associated genes associated with *cis*-eQTLs (*MCM6, DARS*) and four with *trans*-eQTLs (*DGKB, GTF3AP5-AGMO, IL23R/IL12RB2, SLC44A4*). The *DGKB/GTF3AP5-AGMO* region has the most annotation to GWAS hits with variants in the genes showing genome-wide significant associations with T2D^28,29^, fasting plasma glucose traits^30–32^ and glycated hemoglobin^33^. *IL23R/IL12RB2* is a known GWAS locus for age of onset of T2D^34^ while variants in *SLC44A4* have been implicated in the interaction between T2D and iron status biomarkers.^35^ Variants in *MCM6* and *DARS* were recently shown to be associated with total cholesterol change in response to fenofibrate in statin-treated T2D.^36^

Our functional assays in zebrafish focused on the role of the *ZRANB3* ortholog in the pancreas, one of the key tissues in T2D. Knockdown of the gene led to reduced *zranb3* expression and to reduction in pancreatic beta cell number in the developing organism. These observations are consistent with recent evidence suggesting that *Zranb3* is highly expressed in replicating murine beta cells,^37^ suggesting a likely critical role of the gene in production or maintenance of beta-cells. While we found an increased ER stress in the knockdown model, this observation may be due to general stress from loss of the gene given the role of zranb3 as a mediator of DNA replication. These findings in combination with the eQTL findings suggest that the *ZRANB3* locus may act directly, through other loci that it regulates (e.g. *DGKB, GTF3AP5-AGMO, IL23R/IL12RB2*) or in combination with those loci to produce the pathophysiological changes that lead to altered glucose metabolism and T2D.

Integrative analysis of GWAS and transcriptomic studies are increasingly being utilized to identify novel candidate genes which may not have been detected through either type of study alone as illustrated by loci above the null line and towards the upper right quadrant of Figure 5b. We utilized this approach to generate new leads for further studies. We found some candidate genes of which two (*MARCH1* and *LIPC*) are established T2D-related loci. The fact that they were not significant in our study indicates that integrative analysis can boost the capacity of a GWAS to identify and/or prioritize loci for further study. One of the major drawbacks of this type of integrative analysis is the relatively small sample sizes of most transcriptomic datasets^38^, which limits the power of the eQTL studies. However, this limitation would gradually diminish as more data is generated.

The identification of a novel candidate T2D locus in the present study provides further support for the notion that genome analysis studies in diverse global populations have the potential to discover novel risk loci and improve our knowledge of the genetic architecture of many common complex disorders.^39–43^ For T2D, this has been demonstrated in studies which identified *SLC6A11* in Mexicans by the SIGMA Type 2 Diabetes Consortium^44^, *SGCG* in Punjabi Sikhs^45^ and *KCNQ1* in East Asians^46,47^ as novel risk loci for T2D. In the search for novel loci, this strategy of including populations of different ancestries complements the strategy of increasing sample sizes to boost statistical power to detect small effect sizes.

The present study addressed discovery science in the context of under-represented populations in genomic research, partly in response to the lack of diversity and predominance of European ancestry populations in genomic studies.^40,42,48–50^ Several examples now exist for how lack of diversity in genomic studies is resulting in missed opportunities for discoveries and for more robust understanding of heterogeneity in effect sizes across ethnic groups. A recent example from the Population Architecture using Genomics and Epidemiology (PAGE) study demonstrated that one-quarter of genetic associations in the NHGR-EBI GWAS Catalog show significant heterogeneity in effects sizes between ethnicities.^51^ Given that effect sizes are estimates of risk, this implies that risk prediction would vary substantially depending on the ethnic group. It is important to recognize that the effects of this lack of diversity extend beyond discovery science to translational studies because the resulting gaps in knowledge may lead to missed opportunities for developing clinical guidelines, better tailoring of clinical guidelines and treatment protocols and developing new therapeutic agents.^43,52^

The strengths of the present study include a relatively large sample size, a focus on an understudied ancestral group, use of state-of-the-art SNP microarrays and imputation to an African enriched reference panel providing an unparalleled comprehensive opportunity to test millions of common SNPs across African genomes. A potential limitation is that SNPs with small effect sizes are not detectable with the present sample size. More studies in Africans and combined analysis using meta-analytic procedures would overcome this limitation.

In summary, we report a genome-wide analysis of T2D in sub-Saharan Africa in a sample of over 5,000 individuals. The major findings include confirmation of TCF7L2 as genome-wide significant and identification of *ZRANB3* as a novel T2D locus. Functional experiments in zebrafish suggest that ZRANB3 is important in beta cell proliferation and thereby the capacity of the pancreas to respond to insulinogenic stimuli.

## Subjects and Methods

### Ethics statement

All human research was conducted according to the Declaration of Helsinki. The study protocol was approved by the institutional ethics review board of each participating institution. Written informed consent was obtained from each participant prior to enrolment.

### Study participants

Study participants are from the Africa America Diabetes mellitus (AADM) study.^3,6^ This is a study of the genetic epidemiology of T2D in Africa that enrolled participants from Nigeria, Ghana, and Kenya. The study eligibility criteria and enrollment procedures have been described in detail elsewhere.^3,4,6^ Briefly, participants were Africans enrolled through major medical centers in Nigeria (Ibadan, Lagos and Enugu), Ghana (Accra and Kumasi) and Kenya (Eldoret). Participants identified at each center were first consented for the study and underwent the same enrollment procedures, which included collection of demographic details, medical history and clinical examination. Clinic procedures included anthropometry for weight, height, waist circumference and hip circumference; three blood pressure measurements in the sitting position; and collection of fasting blood samples. Weight was measured in light clothes on an electronic scale to the nearest 0.1 kg and height was measured with a stadiometer to the nearest 0.1 cm. Body mass index (BMI) was computed as weight in kg divided by the square of the height in meters.

The definition of T2D was based on the American Diabetes Association (ADA) criteria: a fasting plasma glucose concentration (FPG) ≥ 126 mg/dl (7.0 mmol/l) or a 2-hour post-load value in the oral glucose tolerance test ≥ 200 mg/dl (11.1 mmol/l) on more than one occasion. Alternatively, a diagnosis of T2D was accepted if an individual was on pharmacological treatment for T2D and review of clinical records indicated adequate justification for that therapy. The detection of autoantibodies to glutamic acid decarboxylase (GAD) and/or a fasting C-peptide ≤ 0.03 nmol/l was used to exclude probable cases of type 1 diabetes. Controls were required to have FPG < 110 mg/dl or 2-h post load of < 140 mg/dl and no symptoms suggestive of diabetes (the classical symptoms being polyuria, polydipsia, and unexplained weight loss).

### Biochemistry

Fasting blood samples were assayed for several analytes including glucose, insulin and lipids (total cholesterol, triglycerides, HDL-cholesterol). Fasting plasma glucose was determined using the enzymatic reference method with Hexokinase on COBAS^®^ Analyzer Series(Roche Diagnostics, Indianapolis, IN). Fasting insulin was measured on an Elecsys immunoassay analyzer (Roche Diagnostics) using an electrochemiluminescence technique. Serum lipids were determined enzymatically with COBAS^®^ Analyzer Series (Roche Diagnostics, Indianapolis, IN). The methods have been standardized against the designated CDC reference methods by the manufacturer: CDC reference methods for HDL-cholesterol, Isotope dilution-mass spectrometry (ID-MS) for triglycerides. Serum creatinine and uric acid levels were both measured from fasting blood samples using COBAS^®^ Analyzer Series (Roche Diagnostics, Indianapolis, Indiana).

### Genotyping and Imputation

The 5,231 samples were genotyped on two platforms: 1,808 samples were genotyped using both the Affymetrix Axiom^®^ PANAFR SNP array (an array with ~2.1 million SNPs that is optimized for African ancestry populations (see **Web Resources**) and the Affymetrix 319(R) Exome Array and 3,423 samples were genotyped using the Illumina Consortium array: Multi-Ethnic Global Array (MEGA) (see **Web Resources**). Each of the resulting datasets underwent separate quality control. After technical quality control, sample-level genotype call rate was at least 0.95 for all subjects. Each SNP dataset was filtered for missingness, Hardy-Weinberg equilibrium (HWE) and allele frequency. SNP passing the following filters were retained: missingness < 0.05, HWE p <1 x 10^−6^ and minor allele frequency > 0.01. SNPs that passed quality control were used as the basis for imputation. Imputation of all samples was performed using the African Genome Resources Haplotype Reference Panel (a new African genome reference panel based on 4,956 samples from all African and non-African 1000 Genomes Phase 3 populations and additional African genomes from Uganda, Ethiopia, Egypt, Namibia and South Africa) using the Sanger Imputation Service (see **Web Resources**). The additional African genomes included 2,298 African samples with whole genome sequence data from the African Genome Variation Project (AGVP)^53^ and the Uganda 2,000 Genomes Project (UG2G). This new panel both increased the number of imputed variants and improved the information score and imputation accuracy for African populations when compared with the 1000 Genomes Phase 3 Version 5 reference panel. The resulting imputation dataset of all samples was filtered for variants with minor allele frequency (MAF) >= 0.01 and information score (info) >= 0.3 for association analysis.

### Association analysis

Association analysis was done using the GMMAT (Generalized linear Mixed Model Association Test) R package^54^, a software package for association tests based on generalized linear mixed models. We computed principal components (PCs) using an LD-pruned subset of SNPs (Supplementary Figure 1). Similar to our previous study^4^, we found that the first three PCs were significant and were therefore included in downstream analyses. To account for relatedness between individuals in the sample, we computed a genetic relatedness matrix (GRM) using GEMMA (the Genome-wide Efficient Mixed Model Association algorithm)^55^. Association testing for T2D was done using the mixed logistic model as implemented in GMMAT. This is a score test which was done with the imputed genotype dosages with age, gender, BMI, the GRM and the first three PCs as covariates.

### Statistical power estimates

The power of the study for discovery was estimated using *Quanto*^56^ and assuming an α of 5 × 10^−8^. For a variant with a minor allele frequency (MAF) of 0.05, the study has 80% power to detect a genetic risk ratio (GRR) of 1.7 and 94% power to detect a GRR of 1.8. For a variant with MAF of 0.10, the study has 82% power to detect a GRR of 1.5 and 98% power to detect a GRR of 1.6.

### eQTL annotation of ZRANB3

For eQTL annotation of the genome-wide significant *ZRANB3* SNPs, we utilized data from the Framingham Heart Study (FHS) eQTL Study^38^ accessed via the NCBI Molecular QTL Browser (see **Web Resources**). This is a microarray-based genome-wide study that analyzed both cis and trans eQTLs in whole blood samples from over 5,000 study participants. We chose this database because till date it is the largest, single site study of both *cis*-eQTLs and *trans*-eQTLs. First, we used the haplotype block definition method of Gabriel et al^57^ to construct haplotypes around the two genome-wide significant SNPs, resulting in an 18.7 kb haplotype block around 2:136064024 and a 16.8 kb haplotype bock around 2:136019729. Next, we retrieved significant eQTLs in these two haplotype blocks from the FHS-eQTL Study and identified *cis*-as well as *trans*-eQTL SNP-gene pairs. We then overlaid the gene lists from the retrieved eQTL data on the list of significant associations with T2D in the NHGRI-EBI GWAS Catalog. To provide finer resolution, we annotated the SNPs flanking each genome-wide significant *ZRANB3* SNP for eQTLs.

### Meta-analysis

Given the paucity of genome-wide data on Africans characterized for T2D, we conduct a metaanalysis of our GWAS with an African ancestry dataset: a GWAS of African American samples for T2D (n=8,599) conducted on African American participants from five studies (the Atherosclerosis Risk in Communities (ARIC), the Cleveland Family Study (CFS), the Howard University Family Study (HUFS), Jackson Heart Study (JHS) and Multi-Ethnic Study of Atherosclerosis (MESA) retrieved under controlled access from dbGAP. We used a fixed effects model with inverse weighting of effect sizes as implemented in *METAL*^58^ with double genome inflation correction. As a check, we utilized a meta-analysis method that allows for heterogeneity of effects as implemented in MetaSoft^59^ and obtained essentially the same findings (thus, results from *METAL* are presented). For trans-ethnic meta-analysis, we conduct a meta-analysis for T2D with data from the present GWAS, the African American studies and the DIAGRAM meta-analysis of 13 cohorts imputed from the GoT2D integrated haplotype reference panel^8^.

### Transferability of established type 2 diabetes loci

We looked for evidence of transferability of established T2D loci reported in the literature and curated with the aid of the NHGRI-EBI GWAS Catalog and updated with the latest metaanalysis studies. We considered a p value < 0.05 associated with a SNP with the same direction of effect as evidence of transferability. Where the exact SNP was not present or did not show significant association in our dataset, we examined all SNPs with LD *r*^2^ > 0.3 and within + 250kb of the reported index SNP for association with T2D. Nominal association p values were adjusted for the total number of SNPs within the region using the method of effective degrees of freedom.^60,61^ A locus was considered to show local replication if it had at least one of the tested SNPs with adjusted association p-value < 0.05.

### Zebrafish lines

Experiments were carried out using Tg(*insa*:mCherry).^62^ Adult zebrafish were housed and naturally mated according to standard protocol. All zebrafish work was conducted in accordance with University of Maryland IACUC guidelines.

### Morpholino and CRISPR/Cas9

Morpholino antisense oligonucleotides (MOs) that block splicing (SB) at the splice junction of exon 5 of *zranb3* mRNA were injected into one-to two-cell stage embryos. We designed SB MO (5’-GATACTCCTGCAAAGCAAACAAACA-3’). A control non-specific MO was used (5-CCTCTTACCTCAGTTACAATTTATA-3’). The embryos were grown at 28°C until harvesting for analyses. MO efficacy and off-target toxicity was assessed in cDNA generated from total RNA isolated from homogenates of whole 5 days post fertilization (dpf) larvae and qPCR analysis as per previously reported protocols.^9^

Target sites for CRISPR were determined and designed according to previously published protocols.^9^ We identified target sites within either exon 5 or exon 7 of *zranb3* to which we generated sgRNAs by *in vitro* transcription using the following oligo sequences:

> Exon 5 oligo1:
>
> TAATACGACTCACTATAGGATGGCACGCTTGGCGCTCGTTTTAGAGCTAGAAATAGC
>
> Exon 7 oligo 1:
>
> TAATACGACTCACTATAGGGAATTCGCTGGCGTATTTGTTTTAGAGCTAGAAATAGC
>
> Universal Oligo 2:
>
> AAAAGCACCGACTCGGTGCCACTTTTTCAAGTTGATAACGGACTAGCCTTATTTTAACTTGC
>
> TATTTCTAGCTCTAAAAC

The sgRNAs, at 25 pg/μl, along with Cas9 mRNA, at 300 pg/μl, were microinjected directly into the cell during the single cell stage of embryonic development.

### Zebrafish β-cell analysis

The Tg (*insa*:mCherry) line which labels β-cells specifically by expressing mCherry under the control of the *preproinsulin* (*insa*) promoter was used to quantify the number of β-cells according to previous published protocol.^9^ Briefly, embryos were fixed in 4% PFA, washed in PBST and flat mounted in ProLong^®^Gold antifade (Life Technologies) with the right lateral side facing the coverslip. Sufficient pressure was applied to disrupt the islets in order to visualize individual cells. The number of β-cells was counted manually under an Olympus IX71 fluorescence microscope and imaged and analyzed using CellSens software. The analysis of β-cells was performed on embryos collected from three different injections of either control or test morpholinos.

### Integrative analysis of GWAS with transcriptomic data

To identify potentially novel candidate genes and generate new hypotheses, we leveraged our GWAS to conduct functional gene set based analysis of the AADM GWAS summary statistics using publicly available transcriptomic data on selected T2D-related tissues. Gene-based functional association analytic methods capture the aggregate signals from multiple signals (eQTLs) and may be more appropriate and have more power than testing each variant separately in situations where the expression of a gene is causally related to a trait or disorder and gene expression is regulated by multiple independent eQTLs. We used *EUGENE* v1.3b^63^ using our GWAS summary statistics as input and accounted for LD between variants with the 1000 Genomes AFR data. We restricted our analyses to adipose tissue, skeletal muscle and whole blood using precomputed eQTL reference data provided by the software authors. We used Satterthwaite’s approximation to estimate the significance of the gene-based sum statistic (i.e. *GCTA-fastBAT*)^64^, a method which is more efficient than using simulations to estimate the gene-based sum statistic and is recommended as the default approach to estimate significance.

## Data availability

The GWAS summary statistics are available at dbGap (accession pending); all relevant data are available from the authors upon request under terms consistent with the ethical controls governing the study.

## Acknowledgements

The authors acknowledge with thanks the participants in the AADM project, their families and their physicians. The study was supported in part by the Intramural Research Program of the National Institutes of Health in the Center for Research on Genomics and Global Health (CRGGH). The CRGGH is supported by the National Human Genome Research Institute (NHGRI), the National Institute of Diabetes and Digestive and Kidney Diseases (NIDDK), the Center for Information Technology, and the Office of the Director at the National Institutes of Health (1ZIAHG200362). Support for participant recruitment and initial genetic studies of the AADM study was provided by NIH grant No. 3T37TW00041-03S2 from the Office of Research on Minority Health. Additional support from the NIH includes grants R01DK102001 (N.A.Z.), P30DK072488 (N.A.Z.) and T32DK098107 (T.L.H.).

## Author contributions

CR, AA, GD and FC designed the study; OF, TJ, BE, KA, JA, WB, CA, AA2, JA, DC, CA, GO, JO did participant recruitment, phenotyping and field laboratory assays; AD and SC did molecular laboratory assays and genotyping; AA, AB, JZ, GC, DS did data management and statistical analysis; TH, CL and NZ performed the zebrafish experiments and analyzed the data; AA, AB, AD and NZ drafted the manuscript; CR, DS, AB, AD, NZ and AA edited the manuscript; all authors reviewed and approved the manuscript

## Web resources

Affymetrix Axiom^®^ PANAFR array, https://www.thermofisher.com/order/catalog/product/901788?ICID=search-product ClinVar Accession SCV000191187.1, https://www.ncbi.nlm.nih.gov/clinvar/variation/156804/ dbGaP, https://www.ncbi.nlm.nih.gov/gap Eugene, https://genepi.qimr.edu.au/staff/manuelF/eugene/download.html GMMAT, https://content.sph.harvard.edu/xlin/software.html#gmmat Illumina Multi-Ethnic Global Array (MEGA), https://www.illumina.com/science/consortia/human-consortia/multi-ethnic-genotyping-consortium.html MetaSoft, http://genetics.cs.ucla.edu/meta/index.html The NHGRI-EBI Catalog of published genome-wide association studies, https://www.ebi.ac.uk/gwas/ NCBI Molecular QTL Browser, https://preview.ncbi.nlm.nih.gov/gap/eqtl/studies/ OMIM, http://www.omim.org/ Sanger Imputation Service, https://imputation.sanger.ac.uk/

## References

1 Helgason, A. et al. Refining the impact of TCF7L2 gene variants on type 2 diabetes and adaptive evolution. Nature genetics 39, 218–225, doi:10.1038/ng1960 (2007).

2 Steinthorsdottir, V. et al. A variant in CDKAL1 influences insulin response and risk of type 2 diabetes. Nature genetics 39, 770–775, doi:10.1038/ng2043 (2007).

3 Rotimi, C. N. et al. A genome-wide search for type 2 diabetes susceptibility genes in West Africans: the Africa America Diabetes Mellitus (AADM) Study. Diabetes 53, 838–841 (2004).

4 Adeyemo, A. A. et al. Evaluation of Genome Wide Association Study Associated Type 2 Diabetes Susceptibility Loci in Sub Saharan Africans. Front Genet 6, 335, doi:10.3389/fgene.2015.00335 (2015).

5 Ng, M. C. Y. et al. Meta-Analysis of Genome-Wide Association Studies in African Americans Provides Insights into the Genetic Architecture of Type 2 Diabetes. PLoS Genetics 10:1371/PGENETICS_D_00185R2, doi:10:1371/PGENETICS_D_00185R2 (2014).

6 Rotimi, C. N. et al. In search of susceptibility genes for type 2 diabetes in West Africa: the design and results of the first phase of the AADM study. Annals of epidemiology 11, 51–58 (2001).

7 Chen, G. et al. Common and rare exonic MUC5B variants associated with type 2 diabetes in Han Chinese. PLoS One 12, e0173784, doi:10.1371/journal.pone.0173784 (2017).

8 Fuchsberger, C. et al. The genetic architecture of type 2 diabetes. Nature 536, 41–47, doi:10.1038/nature18642 (2016).

9 Lodh, S., Hostelley, T. L., Leitch, C. C., O’Hare, E. A. & Zaghloul, N. A. Differential effects on beta-cell mass by disruption of Bardet-Biedl syndrome or Alstrom syndrome genes. Hum Mol Genet 25, 57–68, doi:10.1093/hmg/ddv447 (2016).

10 Robu, M. E. et al. p53 activation by knockdown technologies. PLoS Genet 3, e78, doi:10.1371/journal.pgen.0030078 (2007).

11 Sim, X. et al. Transferability of type 2 diabetes implicated loci in multi-ethnic cohorts from Southeast Asia. PLoS Genet 7, e1001363, doi:10.1371/journal.pgen.1001363 (2011).

12 Kraja, A. T. et al. A bivariate genome-wide approach to metabolic syndrome: STAMPEED consortium. Diabetes 60, 1329–1339, doi:10.2337/db10-1011 (2011).

13 Kathiresan, S. et al. Six new loci associated with blood low-density lipoprotein cholesterol, high-density lipoprotein cholesterol or triglycerides in humans. Nat Genet 40, 189–197, doi:10.1038/ng.75 (2008).

14 Waterworth, D. M. et al. Genetic variants influencing circulating lipid levels and risk of coronary artery disease. Arterioscler Thromb Vasc Biol 30, 2264–2276, doi:10.1161/ATVBAHA.109.201020 (2010).

15 Teslovich, T. M. et al. Biological, clinical and population relevance of 95 loci for blood lipids. Nature 466, 707–713, doi:10.1038/nature09270 (2010).

16 Wu, Y. et al. Genetic association with lipids in Filipinos: waist circumference modifies an APOA5 effect on triglyceride levels. J Lipid Res 54, 3198–3205, doi:10.1194/jlr.P042077 (2013).

17 Weissglas-Volkov, D. et al. Genomic study in Mexicans identifies a new locus for triglycerides and refines European lipid loci. J Med Genet 50, 298–308, doi:10.1136/jmedgenet-2012-101461 (2013).

18 Below, J. E. et al. Meta-analysis of lipid-traits in Hispanics identifies novel loci, population-specific effects, and tissue-specific enrichment of eQTLs. Sci Rep 6, 19429, doi:10.1038/srep19429 (2016).

19 Spracklen, C. N. et al. Association analyses of East Asian individuals and transancestry analyses with European individuals reveal new loci associated with cholesterol and triglyceride levels. Hum Mol Genet 26, 1770–1784, doi:10.1093/hmg/ddx062 (2017).

20 Bouhaha, R. et al. TCF7L2 is associated with type 2 diabetes in nonobese individuals from Tunisia. Pathol Biol (Paris) 58, 426–429, doi:10.1016/j.patbio.2009.01.003 (2010).

21 Cauchi, S. et al. European genetic variants associated with type 2 diabetes in North African Arabs. Diabetes & metabolism 38, 316–323, doi:10.1016/j.diabet.2012.02.003 (2012).

22 Danquah, I. et al. The TCF7L2 rs7903146 (T) allele is associated with type 2 diabetes in urban Ghana: a hospital-based case-control study. BMC Med Genet 14, 96, doi:10.1186/1471-2350-14-96 (2013).

23 Guewo-Fokeng, M. et al. Contribution of the TCF7L2 rs7903146 (C/T) gene polymorphism to the susceptibility to type 2 diabetes mellitus in Cameroon. J Diabetes Metab Disord 14, 26, doi:10.1186/s40200-015-0148-z (2015).

24 Nanfa, D. et al. Association between the TCF7L2 rs12255372 (G/T) gene polymorphism and type 2 diabetes mellitus in a Cameroonian population: a pilot study. Clin Transl Med 4, 17, doi:10.1186/s40169-015-0058-1 (2015).

25 Turki, A. et al. Gender-dependent associations of CDKN2A/2B, KCNJ11, POLI, SLC30A8, and TCF7L2 variants with type 2 diabetes in (North African) Tunisian Arabs. Diabetes Res Clin Pract 103, e40–43, doi:10.1016/j.diabres.2013.12.040 (2014).

26 Ng, M. C. et al. Transferability and fine mapping of type 2 diabetes loci in African Americans: the Candidate Gene Association Resource Plus Study. Diabetes 62, 965–976, doi:10.2337/db12-0266 (2013).

27 Ng, M. C. et al. Meta-analysis of genome-wide association studies in african americans provides insights into the genetic architecture of type 2 diabetes. PLoS genetics 10, e1004517, doi:10.1371/journal.pgen.1004517 (2014).

28 Replication, D. I. G. et al. Genome-wide trans-ancestry meta-analysis provides insight into the genetic architecture of type 2 diabetes susceptibility. Nat Genet 46, 234–244, doi:10.1038/ng.2897 (2014).

29 Zhao, W. et al. Identification of new susceptibility loci for type 2 diabetes and shared etiological pathways with coronary heart disease. Nat Genet 49, 1450–1457, doi:10.1038/ng.3943 (2017).

30 Hwang, J. Y. et al. Genome-wide association meta-analysis identifies novel variants associated with fasting plasma glucose in East Asians. Diabetes 64, 291–298, doi:10.2337/db14-0563 (2015).

31 Dupuis, J. et al. New genetic loci implicated in fasting glucose homeostasis and their impact on type 2 diabetes risk. Nat Genet 42, 105–116, doi:10.1038/ng.520 (2010).

32 Manning, A. K. et al. A genome-wide approach accounting for body mass index identifies genetic variants influencing fasting glycemic traits and insulin resistance. Nat Genet 44, 659–669, doi:10.1038/ng.2274 (2012).

33 Wheeler, E. et al. Impact of common genetic determinants of Hemoglobin A1c on type 2 diabetes risk and diagnosis in ancestrally diverse populations: A transethnic genome-wide meta-analysis. PLoS Med 14, e1002383, doi:10.1371/journal.pmed.1002383 (2017).

34 Hamet, P. et al. PROX1 gene CC genotype as a major determinant of early onset of type 2 diabetes in slavic study participants from Action in Diabetes and Vascular Disease: Preterax and Diamicron MR Controlled Evaluation study. J Hypertens 35 Suppl 1, S24–S32, doi:10.1097/HJH.0000000000001241 (2017).

35 Raffield, L. M. et al. Genome-wide association study of iron traits and relation to diabetes in the Hispanic Community Health Study/Study of Latinos (HCHS/SOL): potential genomic intersection of iron and glucose regulation? Hum Mol Genet 26, 1966–1978, doi:10.1093/hmg/ddx082 (2017).

36 Rotroff, D. M. et al. Genetic Variants in HSD17B3, SMAD3, and IPO11 Impact Circulating Lipids in Response to Fenofibrate in Individuals With Type 2 Diabetes. Clin Pharmacol Ther, doi:10.1002/cpt.798 (2017).

37 Klochendler, A. et al. The Genetic Program of Pancreatic beta-Cell Replication In Vivo. Diabetes 65, 2081–2093, doi:10.2337/db16-0003 (2016).

38 Joehanes, R. et al. Integrated genome-wide analysis of expression quantitative trait loci aids interpretation of genomic association studies. Genome Biol 18, 16, doi:10.1186/s13059-016-1142-6 (2017).

39 McCarthy, M. I. Casting a wider net for diabetes susceptibility genes. Nat Genet 40, 1039–1040, doi:10.1038/ng0908-1039 (2008).

40 Need, A. C. & Goldstein, D. B. Next generation disparities in human genomics: concerns and remedies. Trends Genet 25, 489–494, doi:10.1016/j.tig.2009.09.012 (2009).

41 Rosenberg, N. A. et al. Genome-wide association studies in diverse populations. Nat Rev Genet 11, 356–366, doi:10.1038/nrg2760 (2010).

42 Bustamante, C. D., Burchard, E. G. & De la Vega, F. M. Genomics for the world. Nature 475, 163–165, doi:10.1038/475163a (2011).

43 Rotimi, C. N. et al. The genomic landscape of African populations in health and disease. Hum Mol Genet 26, R225–R236, doi:10.1093/hmg/ddx253 (2017).

44 Consortium, S. T. D. et al. Sequence variants in SLC16A11 are a common risk factor for type 2 diabetes in Mexico. Nature 506, 97–101, doi:10.1038/nature12828 (2014).

45 Saxena, R. et al. Genome-wide association study identifies a novel locus contributing to type 2 diabetes susceptibility in Sikhs of Punjabi origin from India. Diabetes 62, 1746–1755, doi:10.2337/db12-1077 (2013).

46 Yasuda, K. et al. Variants in KCNQ1 are associated with susceptibility to type 2 diabetes mellitus. Nature genetics 40, 1092–1097, doi:10.1038/ng.207 (2008).

47 Unoki, H. et al. SNPs in KCNQ1 are associated with susceptibility to type 2 diabetes in East Asian and European populations. Nat Genet 40, 1098–1102, doi:10.1038/ng.208 (2008).

48 Popejoy, A. B. & Fullerton, S. M. Genomics is failing on diversity. Nature 538, 161–164, doi:10.1038/538161a (2016).

49 Bentley, A. R., Callier, S. & Rotimi, C. N. Diversity and inclusion in genomic research: why the uneven progress? J Community Genet 8, 255–266, doi:10.1007/s12687-017-0316-6 (2017).

50 Adeyemo, A. & Rotimi, C. What does genomic medicine mean for diverse populations? Mol Genet Genomic Med 2, 3–6, doi:10.1002/mgg3.63 (2014).

51 Wojcik, G. et al. Genetic diversity turns a new PAGE in our understanding of complex traits. bioRxiv, 188094 (2017).

52 Hindorff, L. A. et al. Prioritizing diversity in human genomics research. Nat Rev Genet, doi:10.1038/nrg.2017.89 (2017).

53 Gurdasani, D. et al. The African Genome Variation Project shapes medical genetics in Africa. Nature 517, 327–332, doi:10.1038/nature13997 (2015).

54 Chen, H. et al. Control for Population Structure and Relatedness for Binary Traits in Genetic Association Studies via Logistic Mixed Models. Am J Hum Genet 98, 653–666, doi:10.1016/j.ajhg.2016.02.012 (2016).

55 Zhou, X. & Stephens, M. Genome-wide efficient mixed-model analysis for association studies. Nat Genet 44, 821–824, doi:10.1038/ng.2310 (2012).

56 Gauderman, W. J. Sample size requirements for association studies of gene-gene interaction. Am J Epidemiol 155, 478–484 (2002).

57 Gabriel, S. B. et al. The structure of haplotype blocks in the human genome. Science 296, 2225–2229, doi:10.1126/science.1069424 (2002).

58 Willer, C. J., Li, Y. & Abecasis, G. R. METAL: fast and efficient meta-analysis of genomewide association scans. Bioinformatics 26, 2190–2191, doi:10.1093/bioinformatics/btq340 (2010).

59 Han, B. & Eskin, E. Random-effects model aimed at discovering associations in metaanalysis of genome-wide association studies. Am J Hum Genet 88, 586–598, doi:10.1016/j.ajhg.2011.04.014 (2011).

60 Bretherton, C. S., Widmann, M., Dymnikov, V. P., Wallace, J. M. & Blade, I. The effective number of spatial degrees of freedom of a time-varying field. J Climate 12, 1990–2009, doi:Doi 10.1175/1520-0442(1999)012<1990:Tenosd>2.0.Co;2 (1999).

61 Ramos, E. et al. Replication of genome-wide association studies (GWAS) loci for fasting plasma glucose in African-Americans. Diabetologia 54, 783–788, doi:10.1007/s00125-010-2002-7 (2011).

62 Pisharath, H., Rhee, J. M., Swanson, M. A., Leach, S. D. & Parsons, M. J. Targeted ablation of beta cells in the embryonic zebrafish pancreas using E. coli nitroreductase. Mech Dev 124, 218–229, doi:10.1016/j.mod.2006.11.005 (2007).

63 Ferreira, M. A. et al. Gene-based analysis of regulatory variants identifies 4 putative novel asthma risk genes related to nucleotide synthesis and signaling. J Allergy Clin Immunol 139, 1148–1157, doi:10.1016/j.jaci.2016.07.017 (2017).

64 Bakshi, A. et al. Fast set-based association analysis using summary data from GWAS identifies novel gene loci for human complex traits. Sci Rep 6, 32894, doi:10.1038/srep32894 (2016).

