## Supplementary Material for "A Novel Type 2 Diabetes Locus in sub-Saharan Africans, *ZRANB3*, is Implicated in Beta Cell Proliferation"

Supplemental Figure S1:  PC plot of genotypes of the AADM participants

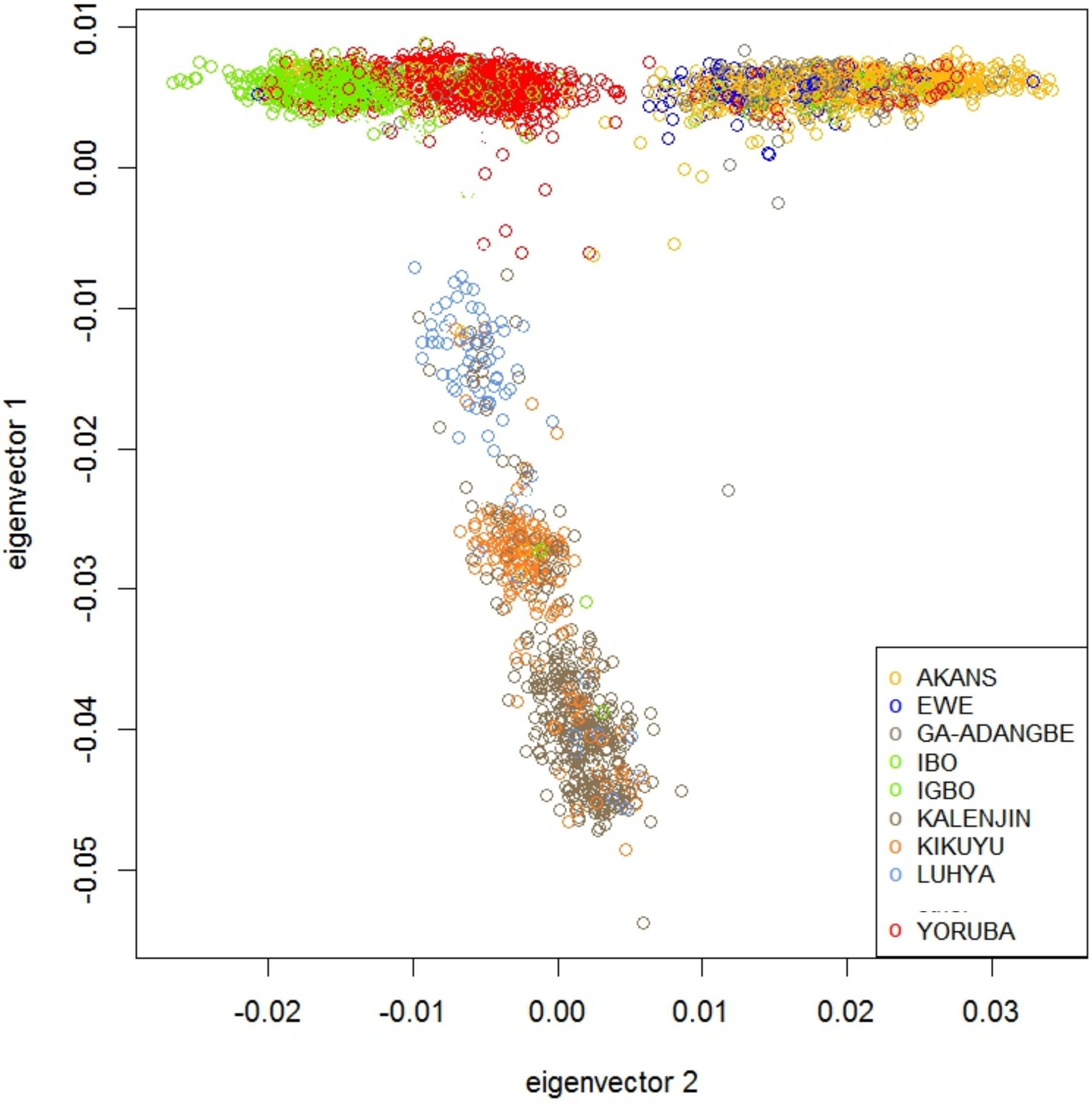

Supplemental Figure S2:  QQ plot for the discovery GWAS for T2D: the AADM Study

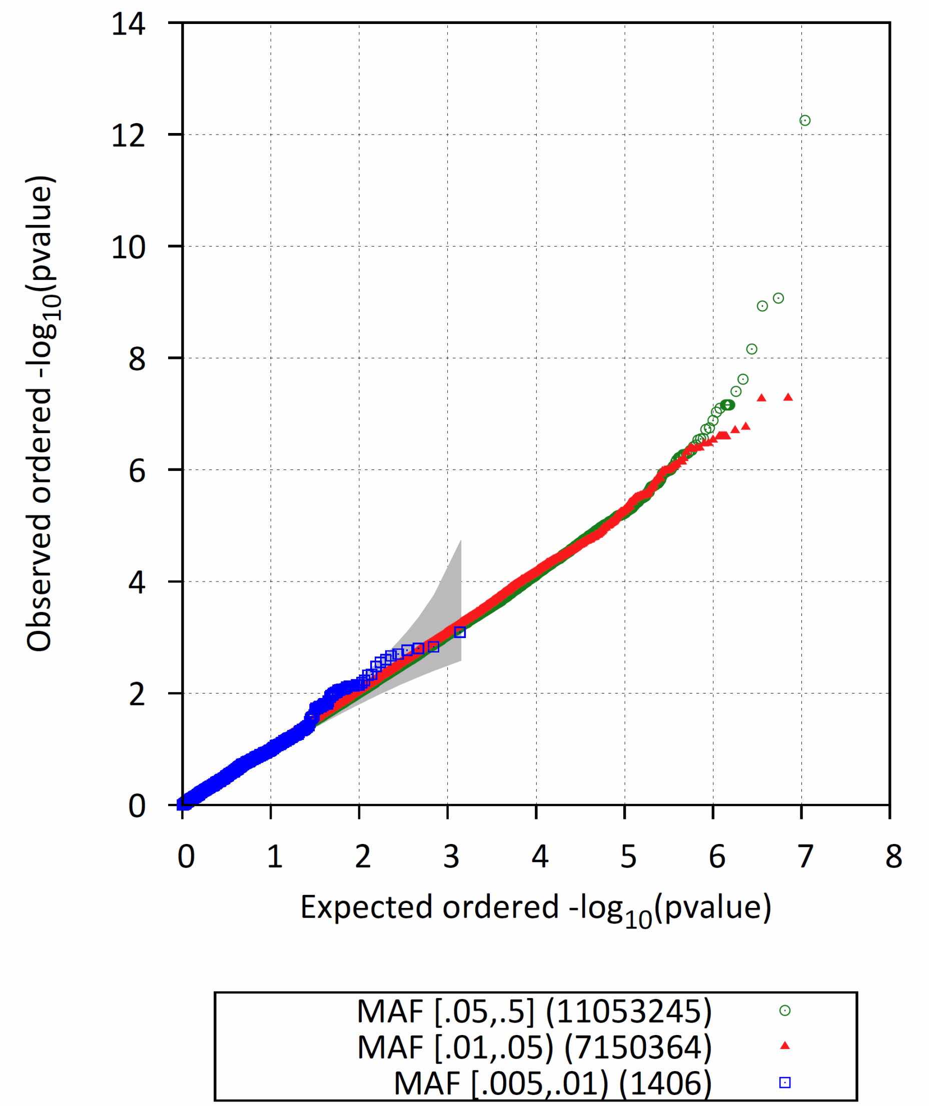

Supplemental Figure S3: ZRANB3 expression in T2D target tissues

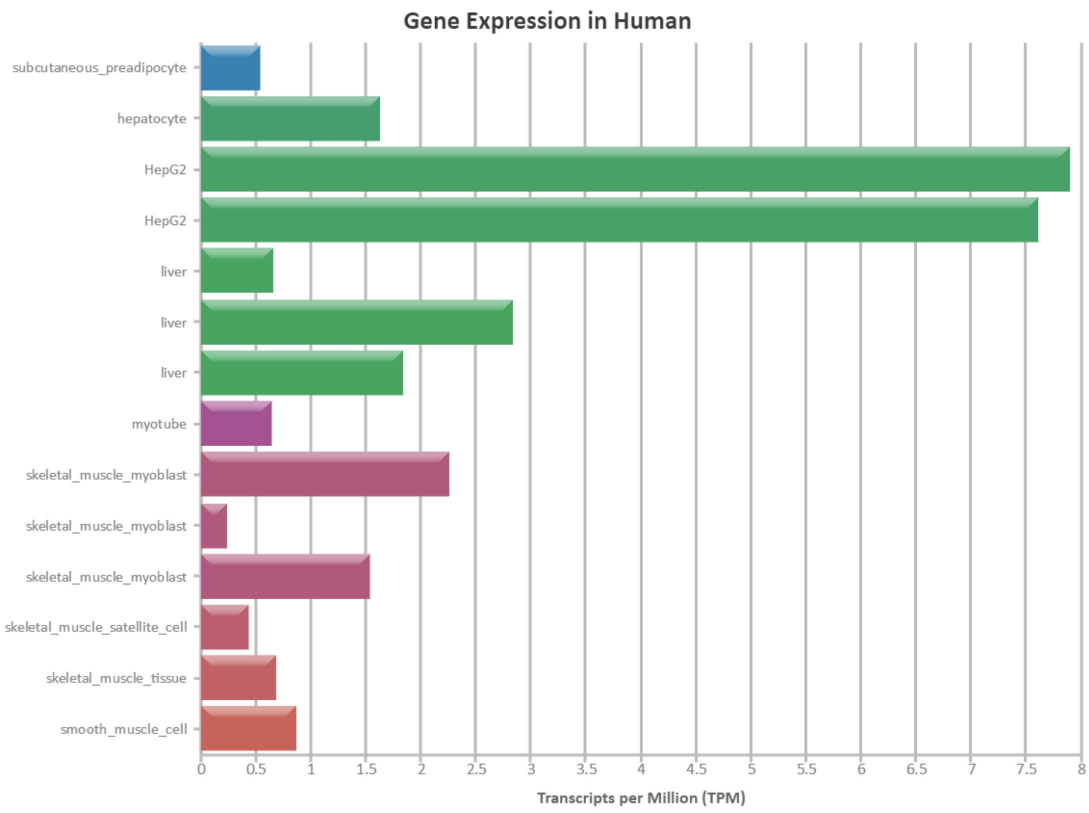


Source: ENCODE

Supplemental Figure S4: Manhattan plot of meta-analysis of African ancestry - AADM and African American - samples

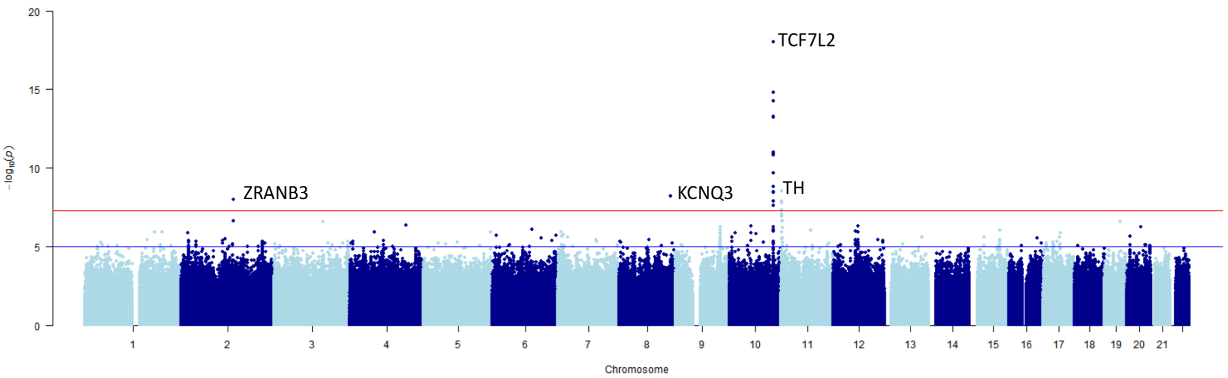


Four genome wide significant loci: *TCF7L2, TH, KCNQ3, ZRANB3*

Supplemental Figure S5:  Evaluation of off-target toxicity in zebrafish knockdonw

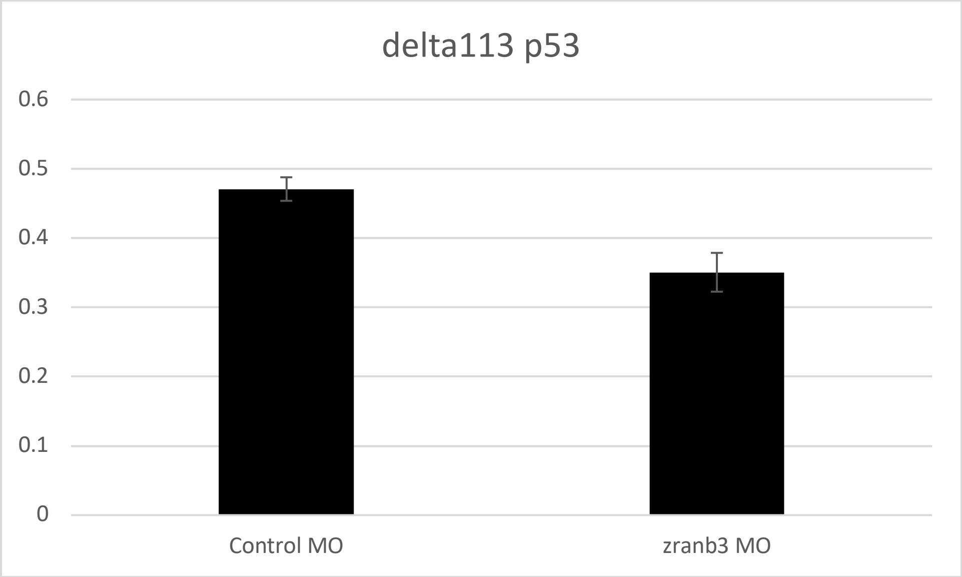


Validation of the efficacy of the *zranb3* MO to significantly suppress *zranb3* mRNA expression without inducing off-target toxicity evaluated using the presence of a marker of MO-induced toxicity, the delta113 isoform of *p53*. Delta113 p53 toxicity data showing no off-target toxicity

Supplemental Figure S6:  Expression of ER stress genes in zranb3 knockdown

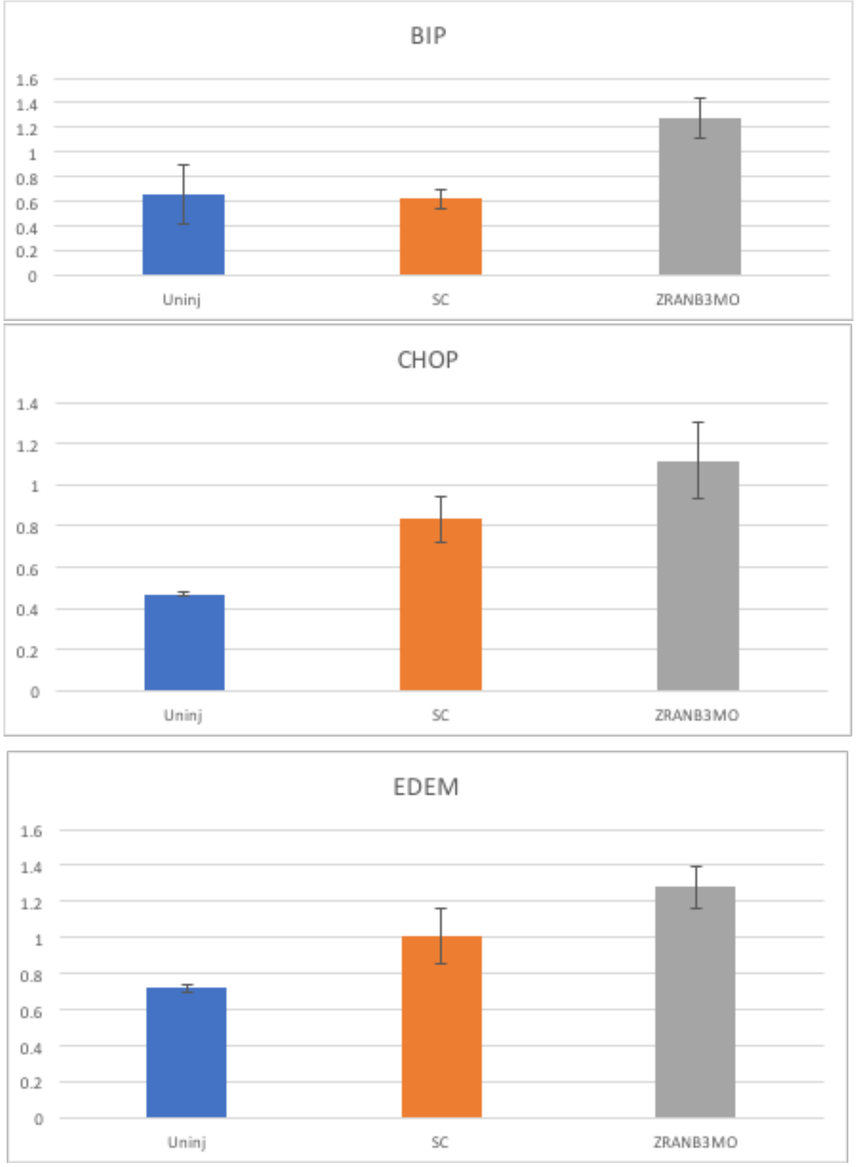

Supplemental Table S1: Local replication of established T2D loci in the AADM Study

| Chr | Pos | Locus | Population | Index SNP | r2 | Best SNP | MAF | BETA | SE | PVAL | df | Adj. P |
|---|---|---|---|---|---|---|---|---|---|---|---|---|
| 1 | 40046093 | MACF1 | European | rs3768321 | 0.5641 | rs111768751 | 0.0138 | 0.536 | 0.2008 | 7.59E-03 | 4.19 | 3.19E-02 |
| 1 | 154320191 | ATP8B2 | Japanese | rs67156297 | 0.3723 | 1:154320191 | 0.0135 | 0.6282 | 0.208 | 2.53E-03 | 8.49 | 2.14E-02 |
| 2 | 43524295 | THADA | European | rs6757251 | 0.6421 | rs534246205 | 0.2919 | -0.1422 | 0.0512 | 5.46E-03 | 8.45 | 4.62E-02 |
| 2 | 57413142 | CCDC85A | Japanese | rs1116357 | 0.3308 | rs56137178 | 0.2368 | 0.1727 | 0.0545 | 1.54E-03 | 6.92 | 1.07E-02 |
| 2 | 161339964 | RBMS1 | non-European | rs1563575 | 0.3083 | rs62175963 | 0.2465 | 0.1669 | 0.0543 | 2.11E-03 | 6.09 | 1.28E-02 |
| 4 | 185721370 | ACSL1 # | Novel 1000G signal | rs60780116 | 0.475 | rs11936062 | 0.6677 | -0.1493 | 0.0488 | 2.24E-03 | 19.37 | 4.33E-02 |
| 6 | 32523888 | HLA-DQA1 # | Novel 1000G signal | rs9271774 | 0.3458 | rs66553215 | 0.6361 | 0.1939 | 0.0545 | 3.73E-04 | 7.8 | 2.92E-03 |
| 6 | 126657472 | CENPW | European | rs11759026 | 0.3055 | rs4897175 | 0.3383 | 0.1634 | 0.0497 | 1.01E-03 | 2.24 | 2.26E-03 |
| 9 | 22289853 | DMRTA1 | Japanese | rs1575972 | 0.9343 | rs12000501 | 0.3117 | -0.2159 | 0.0494 | 1.22E-05 | 13.85 | 1.69E-04 |
| 9 | 84380739 | TLE1 | European | rs9410573 | 0.3095 | rs7033983 | 0.9656 | 0.4575 | 0.1423 | 1.31E-03 | 27.03 | 3.53E-02 |
| 10 | 12246105 | CDC123/ CAMK1D | European | rs11257659 | 0.5412 | rs1320195 | 0.3439 | 0.1687 | 0.0497 | 6.79E-04 | 25.76 | 1.75E-02 |
| 10 | 94281685 | HHEX/IDE | European | rs11187140 | 0.6385 | rs10882074 | 0.0468 | -0.399 | 0.1136 | 4.46E-04 | 9.12 | 4.07E-03 |
| 11 | 2193597 | MIR4686 | Japanese | rs7107784 | 0.539 | rs10770140 | 0.456 | -0.1793 | 0.0473 | 1.48E-04 | 17.18 | 2.54E-03 |
| 11 | 17418477 | KCNJ11 | European | rs5219 | 0.9123 | rs757110 | 0.9168 | -0.2223 | 0.0621 | 3.46E-04 | 10.46 | 3.62E-03 |
| 13 | 80765272 | SPRY2 | European | rs11616380 | 0.3617 | rs2876754 | 0.5826 | 0.1779 | 0.0503 | 4.06E-04 | 15.56 | 6.32E-03 |
| 18 | 57766512 | MC4R | other | rs1942880 | 0.7766 | rs1539952 | 0.2322 | 0.1586 | 0.0531 | 2.85E-03 | 9.78 | 2.79E-02 |

Supplemental Table S2: Top hits from meta-analysis of African ancestry – AADM and 5-cohort African American - studies

| Marker/Alleles/Gene | Weighted N | Zscore | P-value | Direction | Comments |
|---|---|---|---|---|---|
| 10:114758349_C/T_Intron (rs7903146) **TCF7L2** | 14248 | 8.837 | 9.829e-19 | ++ | Expected |
| 11:2187442_C/A_Intron (rs4072825) **TH** | 14248 | 5.934 | 2.96e-09 | ++ | Reported T2D GWAS locus (Imamura *et al* 2016); in the IGF-INS-TH region associated with T2D and metabolic phenotypes; associated with MODY and transient neonatal diabetes; Mutations associated with Segawa syndrome (a neurological syndrome) |
| 8:133465498_G/A_Intron (rs111248619) **KCNQ3** | 14248 | 5.813 | 6.144e-09 | ++ | Not previously known to be associated with T2D; Potassium voltage-gated channel; Associated with neonatal epilepsy |
| 2:136064024_A/T_Intron **ZRANB3** | 3423 | -5.732 | 9.925e-09 | -? | Not previously known to be associated with T2D |
